# Grid-based computational methods for the design of constraint-based parsimonious chemical reaction networks to simulate metabolite production: GridProd

**DOI:** 10.1101/166777

**Authors:** Takeyuki Tamura

## Abstract

Constraint-based metabolic flux analysis of knockout strategies is an efficient method to simulate the production of useful metabolites in microbes. Owing to the recent development of technologies for artificial DNA synthesis, it may become important in the near future to mathematically design minimum metabolic networks to simulate metabolite production. Accordingly, we have developed a computational method where parsimonious metabolic flux distribution is computed for designated constraints on growth and production rates which are represented by grids. When the growth rate of this obtained parsimonious metabolic network is maximized, higher production rates compared to those noted using existing methods are observed for many target metabolites. The set of reactions used in this parsimonious flux distribution consists of reactions included in the original genome scale model iAF1260. The computational experiments show that the grid size affects the obtained production rates. Under the conditions that the growth rate is maximized and the minimum cases of flux variability analysis are considered, the developed method produced more than 90% of metabolites, while the existing methods produced less than 50%. Mathematical explanations using examples are provided to demonstrate potential reasons for the ability of the proposed algorithm to identify design strategies that the existing methods could not identify. The source code is freely available, and is implemented in MATLAB and COBRA toolbox.

**Author summary:** Metabolic networks represent the relationships between biochemical reactions and compounds in living cells. By computationally modifying a given metabolic network of microbes, we can simulate the effect of knockouts and estimate the production of valuable metabolites. A common mathematical model of metabolic networks is the constraint-based flux model. In constraint-based flux balance analysis, a pseudo-steady state is assumed to predict the metabolic profile where the sum of all incoming fluxes is equal to the sum of all outgoing fluxes for each internal metabolite. Based on these constraints, the biomass objective function, written as a linear combination of fluxes, is maximized. In this study, we developed an efficient method for computing the design of minimum metabolic networks by using constraint-based flux balance analysis to simulate the production of useful metabolites.

## Introduction

Finding knockout strategies with minimum sets of genes for the production of valuable metabolites is an important problem in computational biology. Because a significant amount of time and effort is required for knocking out several genes, a smaller number of knockouts is preferred in knockout strategies.

However, the technologies for DNA synthesis are being improved [9]. Although the ability to read DNA is still better than the ability to write DNA, designing synthetic DNA may become important in the near future for the production of metabolites instead of knocking out genes in the original genome. Furthermore, it is more reasonable to design DNA by utilizing already existing genes than to create new genes on a nucleotide level. One to one control relation between each gene and reaction may become possible by modifying existing genes. In contrast to knockout strategies, the number of genes included in the design of synthetic DNA should be as small as possible owing to the requirement of significant experimental effort and time.

Flux balance analysis (**FBA**) is a widely used method for estimating metabolic flux. In FBA, a pseudo-steady sate is assumed where the sum of incoming fluxes is equal to the sum of outgoing fluxes for each internal metabolite [14]. Computationally, FBA is formalized as linear programming (LP) that maximizes biomass production flux, the value of which is called the growth rate (**GR**). The production rate (**PR**) of each metabolite is estimated under the condition that the GR is maximized. Since LP is polynomial-time solvable and there are many efficient solvers, FBA is applicable for use in genome-scale metabolic models. The fluxes calculated by FBA are known to be correspond with experimentally obtained fluxes [24].

Therefore, many computational methods have been developed to identify optimal knockout strategies in genome-scale models using FBA. For example, OptKnock identifies global optimal reaction knockouts with a bi-level linear optimization using mixed integer linear programming (MILP) [1]. The inner problem performs the flux allocation based on the optimization of a particular cellular objective (e.g., maximization of biomass yield, minimization of metabolic adjustment (MOMA [22])). The outer problem then maximizes the target production based on gene/reaction knockouts. RobustKnock maximizes the minimum value of the outer problem [23]. OptOrf and genetic design through multi-objective optimization (GDMO) find gene deletion strategies by MILP with regulatory models and Pareto-optimal solutions, respectively [2, 7]. Dynamic Strain Scanning Optimization (DySScO) integrates the dynamic flux balance analysis (dFBA) method with other strain algorithms [26]. OptStrain and SimOptStrain can identify non-native reactions for target production [8, 16]. In addition to knockouts, OptReg considers flux upregulation and downregulation [17].

Many of the above algorithms are formalized as MILP, which is an NP-hard problem and is computationally very expensive [21]. For example, OptKnock takes around 10 hours to find a triple knockout for acetate production in *E.coli* [11]. To improve runtime performance, different approaches have been developed. OptGene and Genetic Design through Local Search (GDLS) find gene deletion strategies using a genetic algorithm (GA) and local search with multiple search paths, respectively [11, 15]. EMILio and Redirector use iterative linear programs [18, 25]. Genetic Design through Branch and Bound (GDBB) uses a truncated branch and branch algorithm for bi-level optimization [3]. Fast algorithm of knockout screening for target production based on shadow price analysis (FastPros) is an iterative screening approach to discover reaction knockout strategies [13].

Recently, Gu et al. [6] developed IdealKnock, which can identify knockout strategies that achieve a higher target production rate for many metabolites compared to the existing methods. The computational time for IdealKnock is within a few minutes for each target metabolite, and the number of knockouts is not explicitly limited before searching. On the other hand, parsimonious enzyme usage FBA (pFBA) [10] finds a subset of genes and proteins that contribute to the most efficient metabolic network topology under the given growth conditions. Owing to recent development of technologies for artificial DNA synthesis, it may become important in the near future to design minimum metabolic networks that can achieve the overproduction of useful metabolites by selecting a set of reactions or genes from a genome-scale model.

In IdealKnock, ideal-type flux distribution (ITF) and the ideal point=(GR, PR) are important concepts. Since the lower GR tends to result in a higher PR in many cases, IdealKnock uses the minimum “P×TMGR” as the lower bound of the GR and maximizes the PR to find the ITF, where 0 < *P* < 1 and TMGR stands for Theoretical Maximum Growth Rate. Reactions carrying no flux in ITF are treated as candidates for knockout. Although IdealKnock calculates ITF by optimizing the PR with a minimum GR, this method may fail to find the optimal (GR, PR) that achieves a higher PR of target metabolites as discussed in Section.

In this study, we introduce a novel method of calculating parsimonious metabolic networks for producing metabolites (GridProd) by extending the idea of IdealKnock and pFBA. In contrast to IdealKnock, in the calculation of the ideal points, GridProd applies “P” to PR as well as GR. Furthermore, GridProd divides the solution space of FBA into *P*^−2^ small grids, and conducts LP twice for each grid. The area size of each grid is (*P* × *TMGR*) × (*P* × *TMPR*). TMPR stands for theoretical maximum production rate. The first LP obtains reactions included in the designed DNA, and the second LP calculates the PR of the target metabolite under the condition that the GR is maximized for each grid. The design strategy of the grid whose PR is the best is then adopted as the GridProd solution.

Computational experiments were conducted to inspect the efficiency of GridProd using a genome-scale model, iAF1260. The production ability of GridProd strategies was compared to those of IdealKnock and FastPros strategies. GridProd achieves higher PR than IdealKnock for many target metabolites. The average computation time for GridProd is within a few minutes for each target metabolite. The effects of the grid sizes were also inspected. When the solution space was divided into 625 small grids, the obtained PRs were the optimal in the computational experiments, which corresponds to *P*^−1^ = 25.

## Results

### Test for the production of 82 metabolites by exchange reactions

In the first computational experiment, the PRs of the GridProd design strategies were compared to those of the knockout strategies of IdealKnock and FastPros using 82 native metabolites produced by the exchange reactions of iAF1260. For IdealKnock and FastPros, we referred to the results shown in [6].

In the experiments in [6], FastPros took around 3 hours to obtain a strategy for each target metabolite with ten reactions. Therefore, the number of reaction knockouts in that experiments was limited to ten in the experiment of [6]. On the other hand, IdealKnock took 0.3 hours to obtain a strategy for each target metabolite and the knockout number was not limited. All procedures for IdealKnock and FastPros were implemented on a personal computer with 3.40 GHz Intel(R) Core(TM) i7-2600k and 16.0 GB RAM [6].

All procedures for GridProd were implemented on a personal computer with Gurobi, COBRA Toolbox [20] and MATLAB on a Windows machine with Intel(R) Xeon(R) CPU E502630 v2 2.60GHz processors. Although the computers used in the experiments for GridProd and the controls were different, the purpose of this study is not to compare the exact computational times, but rather the of reaction network design each method can find. The results of FastPros may be improved if a larger number of reaction knockouts were allowed.

In the computational experiments described in this study, if the PR was more than or equal to 10^−5^, then the target metabolite was treated as producible. The production ability of each method corresponding to the maximum and minimum PRs calculated by flux variability analysis (FVA) is shown in Table 1. For the maximum case, GridProd produced 75 of the 82 metabolites, while FastPros and IdealKnock produced 45 and 55 metabolites, respectively. For the minimum case, GridProd produced 74 of the 82 metabolites, while FastPros and IdealKnock produced 26 and 40 metabolites, respectively.

**Table 1.**
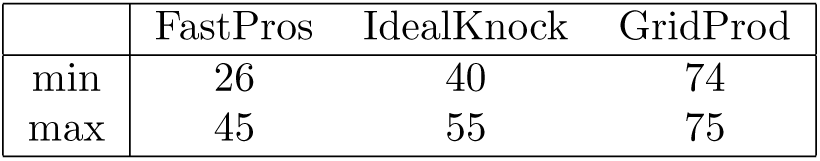
The amount of the 82 iAF1260 target metabolites produced by GridProd, FastPros and IdealKnock strategies. “min” and “max” represent the minimum cases and maximum cases from FVA, respectively.

|  | FastPros | IdealKnock | GridProd |
| --- | --- | --- | --- |
| min | 26 | 40 | 74 |
| max | 45 | 55 | 75 |

The maximum and minimum numbers of reactions used by GridProd for the producible cases were 452 and 406, respectively, for both the maximum and minimum cases from FVA. The average number of reactions used for the producible cases by GridProd were 417.91 and 417.84 for the maximum and minimum cases from FVA, respectively.

The eight target metabolites that were not producible by the GridProd strategies in the minimum cases from FVA are listed in Table 2. The production ability of the eight target metabolites by the FastPros and IdealKnock strategies are also represented in the table. Since IdealKnock could produce seven of the eight target metabolites even for the minimum case from FVA, 81 of the 82 target metabolites were producible by either the GridProd or IdealKnock strategies even for the minimum cases from FVA.

**Table 2.**
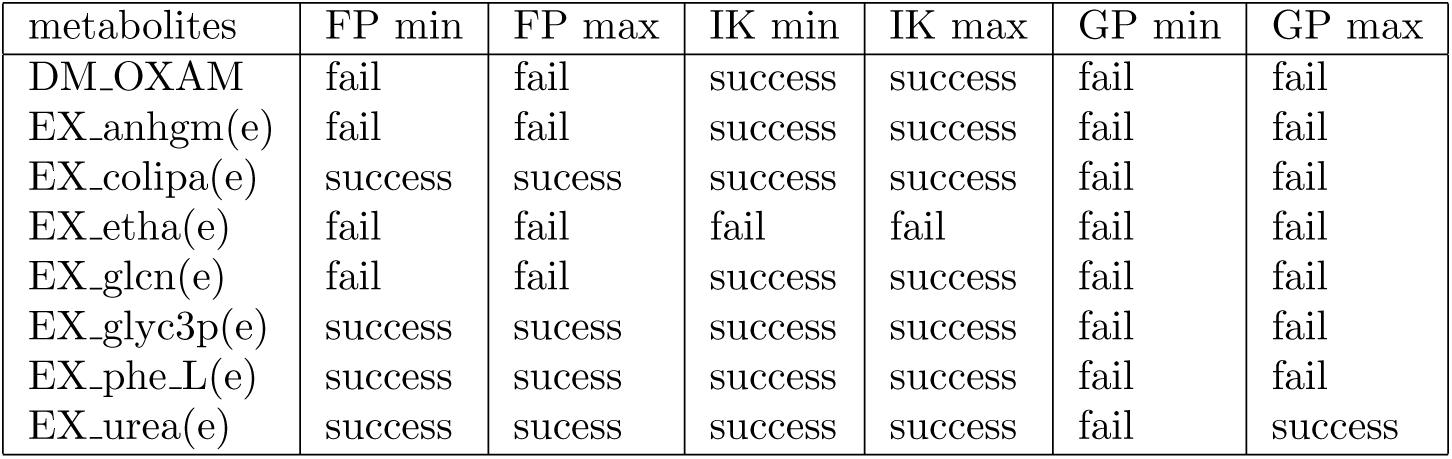
The production ability of each method for the eight target metabolites that were not producible by GridProd in the minimum case from FVA. FP, IK, and GP represent FastPros, IdealKnock and GridProd, respectively. “min” and “max” represent the minimum and the maximum cases from FVA, respectively.

In the second computational experiment, the PRs by the GridProd and IdealKnock strategies were compared for the 82 target metabolites under the condition that the GRs were maximized. As shown in Table 3, for the minimum case from FVA, the PRs of GridProd were higher than those of IdealKnock for 57 of the 82 target metabolites, while the PRs of IdealKnock were higher than those of GridProd for 19 of the 82 target metabolites. The PRs were the same for six target metabolites. As for the maximum case from FVA, the PRs of GridProd were higher than those of IdealKnock for 46 of the 82 target metabolites, while the PRs of IdealKnock were higher than those of GridProd for 35 of the 82 target metabolites. The values were the same for one target metabolite.

**Table 3.** Comparison of the PRs by the GridProd and IdealKnock strategies under the condition that the GRs were maximized. The minimum and maximum cases from FVA were compared, respectively.

|  | GridProd is better | IdealKnock is better | same |
| --- | --- | --- | --- |
| min of FVA | 57 | 19 | 6 |
| max of FVA | 46 | 35 | 1 |

In the third computational experiment, another comparison was conducted between the PRs of GridProd and FastPros under the same condition. The results are shown in Table 4.

**Table 4.** The comparison of the PRs by the strategies of GridProd and FastPros under the condition that the GRs were maximized. The minimum and maximum cases by FVA were compared, respectively.

|  | GridProd is better | FastPros is better | same |
| --- | --- | --- | --- |
| min of FVA | 64 | 11 | 7 |
| max of FVA | 59 | 21 | 2 |

In the fourth computational experiment, various values for P were examined for GridProd. Table 5 shows how many of the 82 target metabolites were produced by the strategies of GridProd for different values of P, where 0 < *P* ≤ 1. When *P* ^−1^ was less than five, the number of producible metabolites was significantly increased as *P*^−1^ became larger. When *P* ^−1^ ≤ 25 held, the number of producible metabolites was almost monotone increase for both the minimum and maximum cases from FVA. When *P*^−1^ = 25 was applied, the numbers of producible metabolites were 74 and 75 for the minimum and maximum cases of FVA, respectively, and this was the best case among the experiments. The average elapsed time for the *P*^−1^ = 25 case was 115.82s.

**Table 5.** The number of producible metabolites by the GridProd strategies in the minimum and maximum cases from FVA for various values of *P*^−1^.

| $P^{-1}$ | min | max | avg elapsed time (s) |
| --- | --- | --- | --- |
| 1 | 1 | 1 | 7.72 |
| 2 | 33 | 35 | 8.97 |
| 3 | 47 | 53 | 9.82 |
| 4 | 58 | 59 | 11.22 |
| 5 | 64 | 64 | 12.09 |
| 6 | 65 | 66 | 14.07 |
| 7 | 65 | 66 | 17.09 |
| 8 | 64 | 64 | 17.55 |
| 9 | 68 | 69 | 21.34 |
| 10 | 71 | 71 | 22.92 |
| 15 | 70 | 71 | 42.57 |
| 20 | 72 | 72 | 77.95 |
| 25 | 74 | 75 | 115.82 |
| 30 | 72 | 72 | 164.78 |
| 100 | 69 | 71 | 1481.84 |

### Test for production of 625 metabolites by transport reactions

In the fifth computational experiment, the PRs by the Grid and FastPros strategies were compared for the 625 target metabolites used in [13]. According to [13], FastPros produced 472 of the 625 metabolites when the number of reaction knockouts was limited to 25, and the average computation time was between 2.6 h and 11.4 h with GNU Linear Programming Kit (GLPK) and MATLAB on a Windows machine with Intel Xeon 2.66 GHz processors.

However, GridProd produced 528 and 535 metabolites for the minimum and maximum cases from FVA, respectively, with *P*^−1^ = 25 as shown in Table 6. Note that the PRs more than or equal to 10^−5^ are treated as producible.

**Table 6.** The number of the 625 target metabolites that were producible by the FastPros and GridProd strategies.

| Method | success | fail | success ratio |
| --- | --- | --- | --- |
| FastPros [13] | 472 | 153 | 75.5% |
| GridProd ($P^{-1} = 25$, min of FVA) | 528 | 97 | 84.5% |
| GridProd ($P^{-1} = 25$, max of FVA) | 535 | 90 | 85.6% |

The PRs of GridProd were better than those of FastPros for 530 of the 625 target metabolites, while FastPros was better than GridProd for 94 target metabolites. They were the same for one metabolite.

For both the minimum and maximum cases from FVA, the maximum, and minimum numbers of reactions used by GridProd for the producible cases were 442 and 404, respectively. The average numbers of reactions used by GridProd for the producible cases were 414.64 and 414.65 for the maximum and minimum cases from FVA, respectively.

## Discussion

FastPros is a shadow price-based iterative knockout screening method. The shadow price in a LP problem is defined as the small change in the objective function associated with the strengthening or relaxing of a particular constraint [13]. Since the knockout candidate is calculated one by one in FastPros, the computational time increases with an increase in the number of knockouts. Therefore, the number of knockouts was limited to less than or equal to 25 in [13]. FastPros showed better performance than OptGene and GDLS for the 625 target metabolites of iAF1260 in the computational experiment described in [13]. When FastPros is combined with OptKnock, improved PRs are observed.

IdealKnock sets the GR to *P* × *TMGR* for various values of *P*, and then maximizes the PRs to obtain the ideal fluxes. All reactions carrying no fluxes in the ideal flux are directly removed. The best results were obtained when *P* was set to 0.05 in [6].

IdealKnock can identify strategies within a few minutes while the number of knockouts is not explicitly limited. For most cases, the sizes of reaction knockout sets were less than 60.

On comparison of the reaction knockout strategies by FastPros and IdealKnock using 82 metabolites based on the computational experiments in [6], IdealKnock exhibited a relatively better performance [6]. FastPros could uniquely predict the overproduction of seven metabolites, while IdealKnock could uniquely predict the production strategies of another 17 metabolites.

While IdealKnock maximizes the PRs with fixed GRs values to find an ideal flux, GridProd imposes the following two constraints

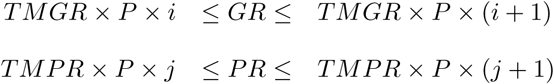

for all integers 1 ≤ *i, j* ≤ *P*^−1^, and then minimizes the sum of absolute values of all fluxes.

The core idea of GridProd is explained using the following examples. Suppose that a toy model of the metabolic network as shown in Fig. 1 is given. {R1,…,R8} and {C1,C2,C3} are sets of reactions and metabolites, respectively. R1 is a source exchange reaction such as glucose or oxygen uptake. R2 is a constant reaction such as ATPM. R7 is the biomass objective function, and R6 is the exchange reaction of the target metabolite. [*a, b*] indicates that *a* and *b* are the lower and upper bounds of the flux for the corresponding reaction. Suppose that the necessary minimum GR is 1 in this example.

**Figure 1.**
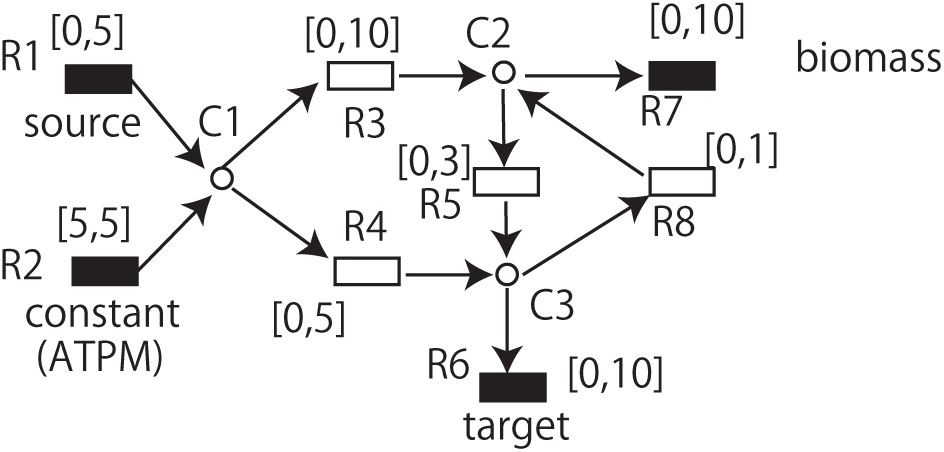
A toy example of the metabolic network, in which GridProd can identify the optimal strategy but IdealKnock cannot under the condition that GR is maximized.

In the original state, if GR is maximized, GR becomes 10 by (R1,R2,R3,R7) = (5,5,10,10). However, PR becomes 0 since the sum of upper bounds of R1 and R2 is 10, and all flow from R1 and R2 goes to R7. If PR is maximized, R6 becomes 8 since R4=5 and R5=3 are the bottle necks. Therefore, TMGR and TMPR are 10 and 8, respectively. If PR is maximized for a fixed GR as in IdealKnock, PR becomes max(10-GR,8).

The optimal design strategy in this network to obtain the maximum PR under the condition that GR is maximized is to knockout R3 where R5 is optional. In this case, (GR,PR)=(1,4) is obtained. Note that the minimum necessary GR is set to 1 in this example. If R3 is not knocked out, (GR,PR)=(10,0) is always obtained.

Suppose we adopt the strategy where a set of reactions not included in the initially obtained flux is knocked out. If GR > 1 is fixed and PR is maximized, R3 must be used since the upper bound of R8 is 1. Therefore, R3 is not knocked out, and then (GR,PR)=(10,0) is obtained when GR is maximized. Next, suppose that *GR* ≤ 1 is fixed and PR is maximized. Note that setting *GR* < 1 is possible for the first LP, although the necessary minimum GR is 1 for the second LP. Then, (R3,R5)=(3+GR,5) is obtained, and PR is 8. Since R3 is not knocked out in this case, (GR,PR)=(10,0) is obtained when GR is maximized. Thus, the ideal flow-based approach that maximizes PR for the fixed values of GR cannot identify the strategy of knocking out R3 and does not obtain PR=4.

To address this, GridProd applies P to both GR and PR. However, there may be multiple flows that satisfy the given constraints for GR and PR. For example, if (GR,PR)=(1,4) is given as the constraints, there are multiple flows satisfying these constraints. However, R4 must be used in any flow since the upper bound of R5 is 3. If R4 is 5, then R8 is 1 and R3=R5=0 holds. If R3 and R5 are knocked out, (GR,PR)=(1,4) is achieved. However, if R4< 5 holds, then R3 and R8 must be used and R5 is optional. Then (GR,PR)=(10,0) is obtained. Since GridProd minimizes the total sum of absolute values of fluxes, (GR,PR)=(1,4) is obtained by knocking out R3.

To discuss the effects of the size of each grid, we analyze each case where GR∈ {0, 1, 2} and PR∈ {3, 4, 5} are given in the following. Suppose that (GR,PR)=(1,5) or (GR,PR)=(2,4) is given. Then, R4 must be used since the upper bound of R5 is 3. In addition to R4, R3 also must be used since *R*1 + *R*2 = 6 must hold. R5 and R8 are optional. In every case, the consequent reaction knockout results in (GR,PR)=(10,0). Note that the necessary minimum growth is assumed as 1 in this example, however, GR is allowed to be less than 1 if GR ≥ 1 is satisfied in the consequent strategies. When (GR,PR)=(0,5) is given, R4 must be used since the upper bound of R5 is 3. R3 is optional. If R3 is used, then R5 must be used, and R8 is optional. If {R3,R5,R8} is knocked out, then GR becomes 0 and minGrowth cannot be satisfied. If only R8 is knocked out, then (GR,PR)=(10,0) is obtained. When (GR,PR)=(2,3) is given, there are multiple flows. If R4 is not used, then R3 and R5 must be 5 and 3, respectively. Consequently, R4 and R8 are knocked out, and then (GR,PR)=(10,0) is obtained. If R4 is used, R3 must be used since the upper bound of R8 is 1. R5 and R8 are optional. Then, (GR,PR)=(10,0) is obtained. When (GR,PR)=(2,5) is given, R4 must be used since the upper bound of R5 is 3. Since the upper bound of R8 is 1, R3 must be used. R5 and R8 are optional. Then, (GR,PR)=(10,0) is obtained. If (GR,PR) is (0,3), (1,3), or (0,4), then there is no flux satisfying the condition since the lower bound of R2 is 5.

Therefore, when GR∈{0, 1, 2} and PR∈ {3, 4, 5} are given for the first LP, the consequent (GR,PR) obtained by the second LP is represented in Table 7. Although (GR,PR) is given as exact values in the above example for simplicity, they are given as constraints represented by the inequalities in GridProd. Suppose that the size of each grid is relatively large, and the corresponding constraints are 0 ≤ *GR* ≤ 2 and 3 ≤ *PR* ≤ 5. Then, one of the possible obtained flow by the first LP is (R1,…,R8)=(0,5,0,5,0,0,0,0) since the sum of absolute values of fluxes are minimized in the first LP of GridProd. Consequently, R3, R5, and R8 are knockedout. Then the second LP is not feasible. However, if the size of each grid is small and the corresponding constraints are 1 – *∊* ≤ *GR* ≤ 1 + *∊* and 4 – *∊* ≤ *PR* ≤ 4 + *∊* where *∊* is a small positive constant, then (GR,PR)=(1,4) is achieved in the second LP. Therefore, the size of each grid affects the resulting PR of the target metabolites. Table 5 shows that as *P*^−1^ becomes larger, the production ability improves when *P*^−1^ ≤ 25. However, when *P*^−1^ > 25 holds, the production ability does not improve as *P*^−1^ becomes larger. This indicates that the necessary minimum size of *∊* in the above example is related to the necessary minimum size of *P*^−1^.

**Table 7.**
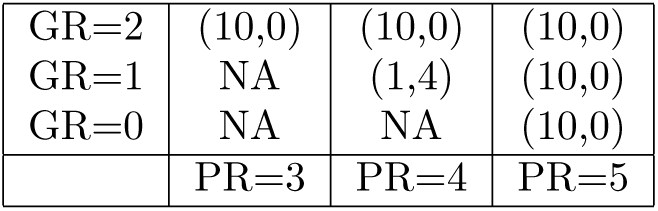
Values of (GR,PR) obtained by the second LP of GridProd when GR∈ {0, 1, 2} and PR∈ {3, 4, 5} are given as the constraints for the first LP.

Table 1 shows that GridProd could find the strategies for producing at least 20 target metabolites that IdealKnock could not identify. Potential reasons for this improvement include the effects of the parsimonious-based approach and the grid-based approach as explained above. Since 74 of the 82 target metabolites were producible via the GridProd strategies even for the minimum cases from FVA, there are eight target metabolites that may not be producible by the GridProd strategies. Table 2 shows that FastPros and IdealKnock produced many of these eight target metabolites. Since IdealKnock could produce all target metabolites but ‘Ex_etha(e)’ even for the minimum cases from FVA, 81 of the 82 target metabolites were producible either by FastPros, IdealKnock or GridProd. The reason as to why none of the methods could identify a strategy to produce ‘Ex_etha(e)’ requires further investigation.

GridProd computes the design of chemical reaction networks by choosing reactions used in the first LP. Because many reactions in iAF1260 are not associated with genes, it is not directly possible to extend the idea of GridProd for the selection of a set of genes.

### Materials and Methods

The pseudo-code of GridProd is as follows.

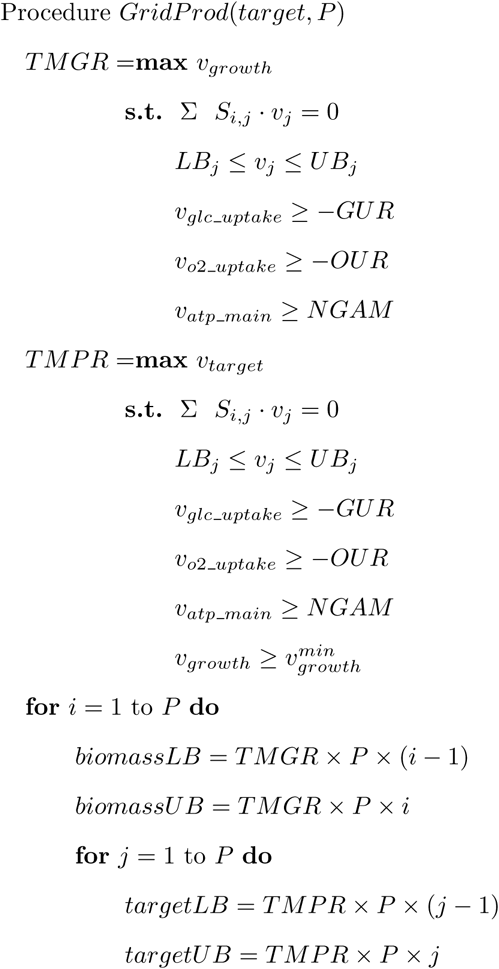

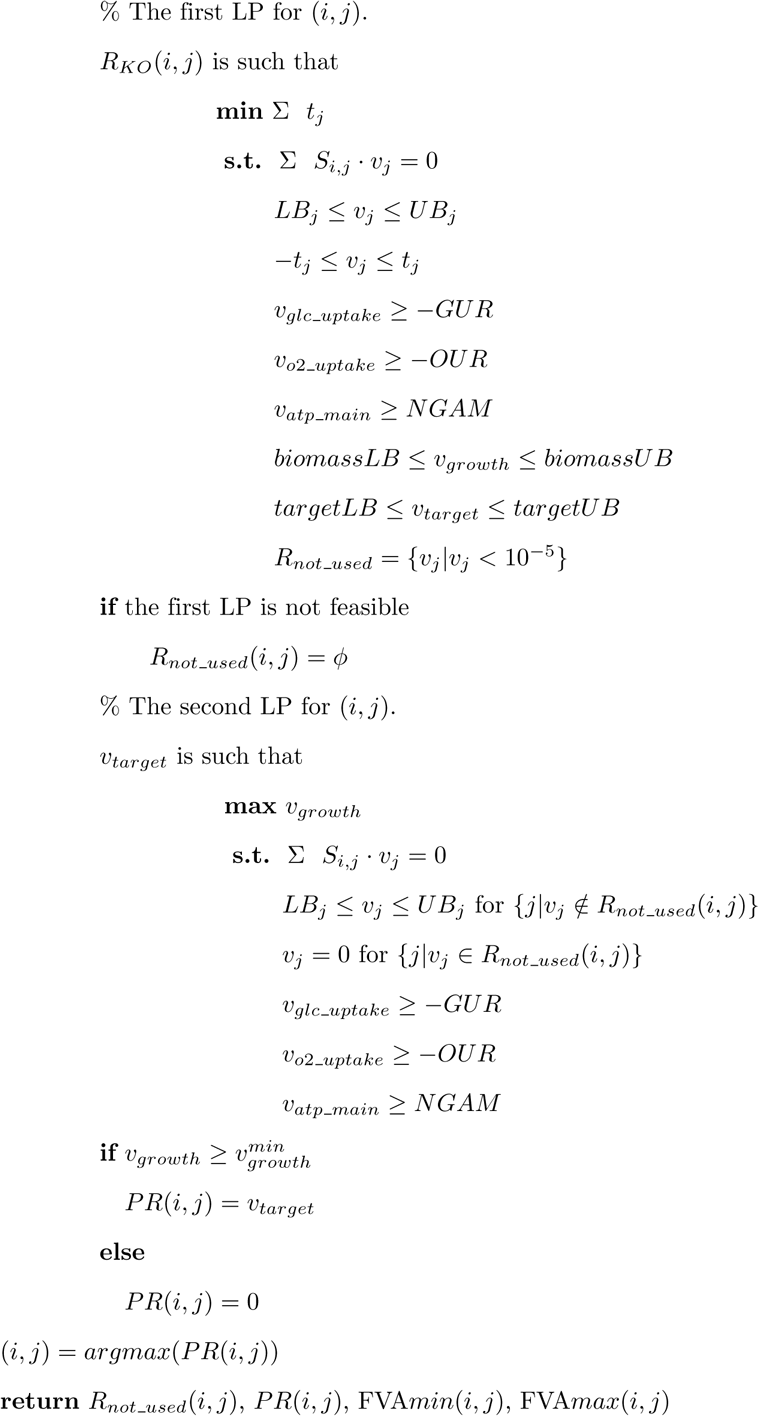

In the above pseudo-code, the TMGR and TMPR are calculated first. *S_i,j_* is the stoichiometric matrix. *LB_j_* and *UB_j_* are the lower and upper bounds of *v_j_*, respectively, that represents the flux of the *j*th reaction.

*v_glc_uptake_*, *v*_*o*2_*uptake*_, and *v_atp_main_* are the lower bounds for the uptake rate of glucose (GUR), the oxygen uptake rate (OUR), and the non-growth-associated APR maintenance requirement (NGAM), respectively. 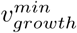 is the minimum cell growth rate.

In each grid, LP is conducted twice. “*biomassLB*” and “*biomassUB*” represent the lower and upper bounds of GR, respectively. Similarly, “*targetLB*” and “*targetUB*” represent the lower and upper bounds of PR, respectively, which are used as the constraints in the first LP. Each grid is represented by the two constraints, “*biomassLB* ≤ *v_growth_* ≤ *biomassUB*” and “*targetLB* ≤ *v_target_* ≤ *targetUB*”. *TMPR* × *P* and *TMGR* × *P* represent the horizontal and vertical lengths of the grids, respectively.

In the solution of the first LP, a set of reactions whose fluxes are almost 0 (less than 10^−5^) are represented as *R_not_used_*, which is used as a set of unused reactions in the second LP. In the second LP, none of the “*biomassLB*”, “*biomassUB*‘”, “*targetLB*”, and “*targetUB*” are used, but the fluxes of the reactions included in *R_not_used_* were forced to be 0. If the obtained PR is more than or equal to 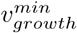 in the solution of the second LP, the value of PR is stored to *PR*(*i, j*). Otherwise 0 is stored. Finally, the (*i, j*) that yields the maximum value in *PR*(*i, j*) is searched, and the corresponding *R_not_used_*(*i, j*) and *PR*(*i, j*) are obtained. The minimum and maximum PRs from FVA for *R_not_used_*(*i, j*) are also calculated. 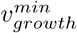 is set to 0.05 in GridProd as in [13].

### Genome-scale metabolic model of *Escherichia coli*

iAF1260 is a genome-scale reconstruction of the metabolic network in *Escherichia coli K-12* MG1655 and includes 1260 open reading frames and more than 2000 transport and intracellular reactions [5]. We used iAF1260 as an original mathematical model of metabolic networks. To simulate the production potential for each target metabolite in this model, we added a transport reaction for the target metabolite if it were absent in the original model, which was assumed to be a diffusion transport as in [13].

In our computational experiments, glucose was the sole carbon source, and the GUR was set to 10 mmol/gDW/h, the OUR was set to 5 mmol/gDW/h, the NGAM was set to 8.39, and the minimum cell growth rate 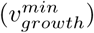 was set to 0.05, as in [13]. These conditions correspond to microaerobic conditions, where the oxygen uptake is insufficient to oxidize all NADH produced in glycolysis and the tricarboxylic acid cycle in the electron transfer system. This relatively low OUR was chosen because higher production yields of target metabolites can be obtained under such conditions compared with under the higher OUR when carbon is mainly used to generate biomass and CO_2_ [13]. Other external metabolites such as CO_2_ and NH_3_ were allowed to be freely transported through the cell membrane in accordance with [5]. Although it is not realistic to assume that large molecules diffuse out of *E. coli*, it may become important in the near future to compute the design of parsimonious chemical reaction networks to produce various metabolites.

For constraint-based analysis using GSMs, simplified models are often considered to reduce computational time [4, 19]; such models provide identical flux estimation and screening results as the original model [12]. However, in this study, we used the original iAF1260 model as opposed to such simplified models because it takes only a few minutes for GridProd to obtain a solution for each target metabolite in most cases.

## Supporting information

**S1 File** All source codes and the solutions obtained by GridProd in the computational experiments described in this manuscript are included.

## Acknowledgments

*Funding:* T.T. was partially supported by grants from JSPS, KAKENHI #16K00391 and #16H02485.

*Conflict of interests:* none declared.

